# HamHeat: A fast and simple package for calculating Hamming distance from multiple sequence data for heatmap visualization

**DOI:** 10.1101/2020.03.26.009258

**Authors:** Alexey V. Rakov, Dieter M. Schifferli, Shu-Lin Liu, Emilio Mastriani

## Abstract

The problem of fast calculation of Hamming distance inferred from many sequence datasets is still not a trivial task. Here, we present HamHeat, as a new software package to efficiently calculate Hamming distance for hundreds of aligned protein or DNA sequences of a large number of residues or nucleotides, respectively. HamHeat uses a unique algorithm with many advantages, including its ease of use and the execution of fast runs for large amounts of data. The package consists of three consecutive modules. In the first module, the software ranks the sequences from the most to the least frequent variant. The second module uses the most common variant as the reference sequence to calculate the Hamming distance of each additional sequence based on the number of residue or nucleotide changes. A final module formats all the results in a comprehensive table that displays the sequence ranks and Hamming distances.

**Availability and implementation:** HamHeat is based on Python 3 and AWK, runs on Linux system and is available under the MIT License at: https://github.com/alexeyrakov/HamHeat.

## 1. Introduction

The calculation of pairwise Hamming distances is one of the most common tasks in string metrics to measure the edit distance between two sequences [Pinheiro et al., 2012]. There are numerous algorithms to determine Hamming distances [Pappalardo et al., 2009; Zhang et al., 2013; Kopelowitz et al., 2018] and some of them are implemented as functions in programming languages (e.g. hamming_distance(x, y) in Python 2.3+ or hamming.distance(x, y) in R). However, all these methods compare only two sequences. In this article, we present a straightforward and practical package for calculating the Hamming distance from large sets of aligned protein or DNA sequences of same lengths. The algorithm automatically compares the most frequent variant with all the other variants for ranking purposes and calculates all the Hamming distances based on the amino acid residue or nucleotide changes. The data are presented in the form of a table that can serve as input for further visualizing software. HamHeat is available as open-source code from our GitHub repository under the MIT License.

## 2. Implementation

Briefly, HamHeat implements the Hamming distance calculation in three modular steps. On the first step, the program ranks data from the most common variant to the least one. On the second step, variants are ranked by using the most common variant as the reference sequence to calculate the number of changes. A final module presents the results in the form of a comprehensive table.

The three modules of HamHeat are written in Python and run consecutively from the command line using the single bash script called *hamheat.sh*. The process can be visualized as shown in Fig. 1.

**Fig. 1.**
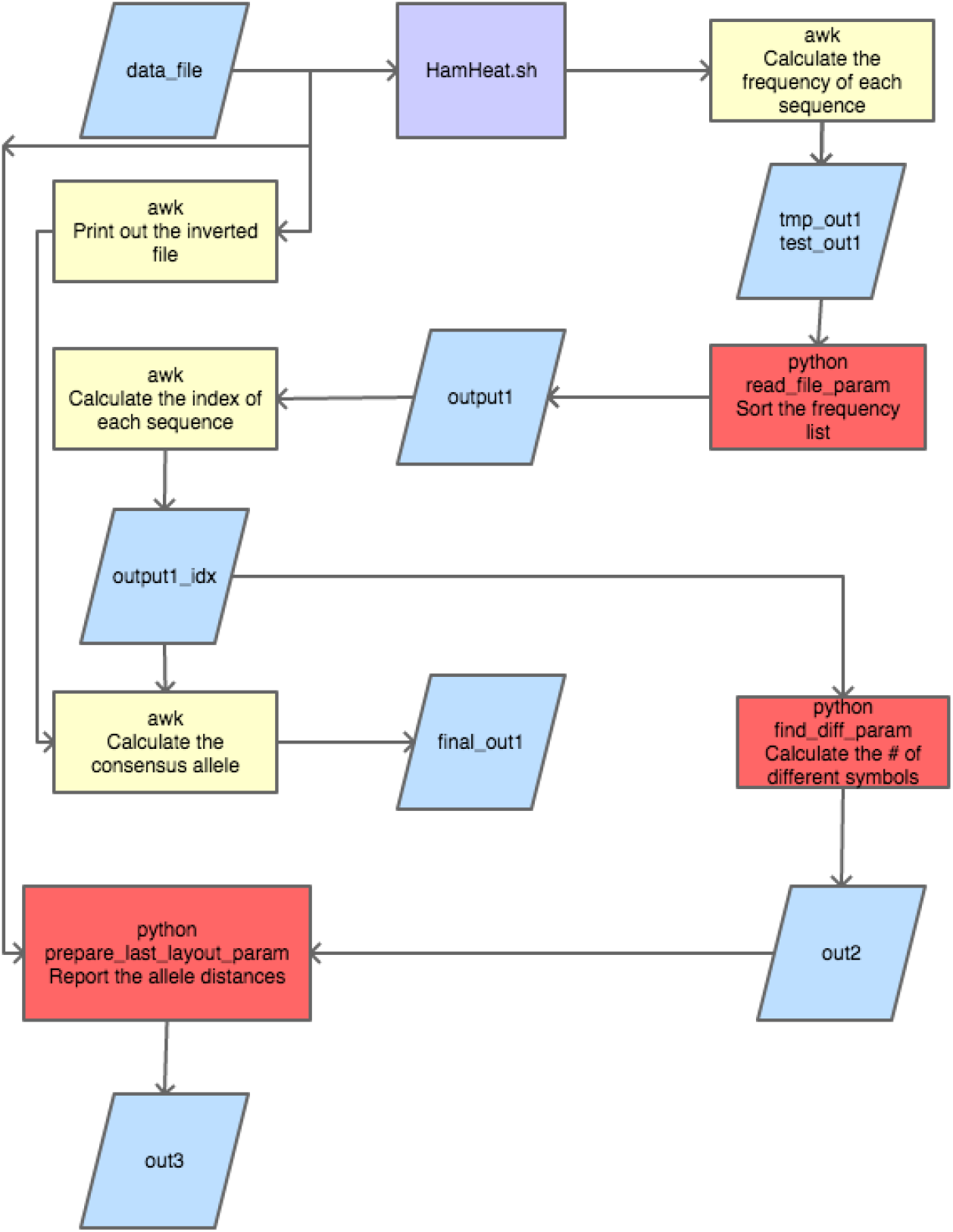
HamHeat process flowchart. The chart shows the process information flow, with the input/output files (blue boxes), the “wrap” bash script (purple box), the AWK commands (yellow boxes), and the Python modules (red boxes).

Users only need to specify the name of an input file in tab-delimited format with two columns, with the first column for the sequence names and the second column for the aligned sequences. There are no limitations for the number and length of the aligned sequences. The only condition is that all the sequences must have the same length. The output file will also be in a tab-delimited layout and composed of three columns, consisting of the aligned sequences, the name of sequences, and the Hamming distances, where a zero distance (0) is given for the most frequent variant which is selected as the reference sequence by HamHeat.

The resulting tab-delimited file may then be used as input file for any of heatmap producing software. For instance, we used the Morpheus matrix visualization software (https://software.broadinstitute.org/morpheus/) in our previously published study [Rakov et al., 2019].

## 3. Results

We successfully tested our script in the Linux Ubuntu environment. The script was applied to the real *Salmonella* allelic data of our recent publication [Rakov et al., 2019]. Figure S1 (additional file 5) from this publication shows the heatmap for 70 virulence factors alleles for 500 *Salmonella* genomes. For this figure, the HamHeat results for each of the 70 alleles were combined in one file to be used as the input file for the Morpheus matrix visualization software.

## 4. Conclusions

HamHeat enables fast Hamming distance calculations for multiple aligned sequence data that can be further used for heatmap visualizing software. This is, to our knowledge, the first user-oriented implementation of the Hamming distance calculation for multiple sequence data and for heatmap visualization. We believe that HamHeat will be particularly useful for allelic variation projects.

## 5. Availability of data

Project name: HamHeat

Project home page: https://github.com/alexeyrakov/HamHeat

## 8. Authors’ contributions

AVR conceived of the presented idea, designed and wrote the software, performed the analyses, and wrote the manuscript. DMS and SLL co-wrote the manuscript. EM designed and wrote the software, drawn a figure, and co-wrote the manuscript. All authors discussed the results and contributed to the final manuscript.

## 9. Acknowledgements

We thank the developers of the Morpheus matrix visualization and analysis software from the Broad Institute of MIT and Harvard.

